# Acute toxicity effects of pesticides on beneficial organisms – Dispelling myths for a more sustainable use of chemicals in agricultural environments

**DOI:** 10.1101/2024.01.25.577310

**Authors:** Luis Mata, Rosemary A. Knapp, Robert McDougall, Kathy Overton, Ary A. Hoffmann, Paul A. Umina

## Abstract

Agricultural practitioners, researchers and policymakers are increasingly advocating for integrated pest management (IPM) to reduce pesticide use while preserving crop productivity and profitability. Selective pesticides, putatively designed to act on pests while minimising impacts on off-target organisms, have emerged as one such option – yet evidence of whether these compounds control pests without adversely affecting natural enemies and other beneficial species (henceforth beneficials) remains scarce. At present, the selection of pesticides compatible with IPM often considers a single (or a limited number of) widely distributed beneficial species, without considering undesired effects on co-occurring beneficials. In this study, we conducted standardised laboratory bioassays to assess the acute toxicity effects of 20 chemicals on 15 beneficial species at multiple exposure timepoints, with the specific aims to: (1) identify common and diverging patterns in acute toxicity responses of tested beneficials; (2) determine if the effect of pesticides on beetles, wasps and mites is consistent across species within these groups; and (3) assess the impact of mortality assessment timepoints on International Organisation for Biological Control (IOBC) toxicity classifications. Our work demonstrates that in most cases, chemical toxicities cannot be generalised across a range of beneficial insects and mites providing biological control, a finding that was found even when comparing impacts among closely related species of beetles, wasps and mites. Additionally, we show that toxicity impacts increase with exposure length, pointing to limitations of IOBC protocols. This work challenges the notion that chemical toxicities can be adequately tested on a limited number of ‘representative’ species; instead it highlights the need for careful consideration and testing on a range of regionally and seasonally relevant beneficial species.

## 1. Introduction

Pesticide pollution in the air, on land, and in the water is a global phenomenon responsible for a wide array of negative effects on people and other species across both natural and anthropogenic environments (Köhler and Triebskorn 2013; Hallmann et al. 2014; Jacobsen & Hjelmsø 2014; Eng et al. 2019; Sharma et al. 2019; Tang et al. 2021; de Montaigu and Goulson 2023; Grondona et al. 2023; Maggi et al. 2023). Pesticides are particularly detrimental to insects, arachnids and other invertebrates, being one of the most acute drivers of concerning population declines (van Lexmond et al. 2015; Forister et al. 2019; Eggleton 2020; Wagner 2020; Wagner et al. 2021). Counterintuitively however, a gamut of pesticides engineered specifically to target invertebrate species are remarkably effective at delivering a diverse array of health and economic benefits to people – a incongruency that has been aptly termed ‘The Pesticide Paradox’ (Enserink et al. 2013).

In agricultural environments, insecticides and miticides (herein referred to as pesticides) remain the dominant management option used by food and textile producers to control a large diversity of invertebrate pests (Cooper and Dobson 2007; Schreinemachers & Tipraqsa 2012; Dewar and Denholm 2017). Their widespread and indiscriminate use, however, has adversely impacted agroecosystems across the world, resulting in a wide array of detrimental outcomes, including: (1) acute toxicity, sub-lethal and transgenerational effects on non-target species (Haynes 1988; Boatman et al. 2004; Schulz et al. 2021; Zioga et al. 2023), including natural enemies and other beneficial species (Desneux et al. 2007; Frampton and Dorner 2007; Mills et al. 2016; Siviter & Muth 2020; Wu et al. 2022; McDougall et al. in press); (2) emergence of secondary pest outbreaks (Hill et al. 2017; Buitenhuis et al. 2023); and (3) the evolution of pesticide resistance (Foster et al. 2017; Wang et al. 2020; Umina et al. 2019; Arthur et al. 2021; Umina et al. 2022). Not surprisingly, farmers, agronomy advisers, researchers, and policymakers across grains, horticulture, and textile industries, are showing a growing interest – and in many instances openly advocating for – more sustainable pest control management options based on reducing pesticide use while preserving crop productivity and profitability (Lechenet et al. 2017; Wilson and Tisdell 2001; Tilman et al. 2002; Foley et al. 2011; Hossard et al. 2014; Dudley et al. 2017; Lykogianni et al. 2021; Schneider et al. 2023).

Integrated Pest Management (IPM) is a well-established approach to pest control (Kogan 1998; Deguine et al. 2021), in which preference is given to non-chemical over chemical methods, assuming they provide the necessary pest suppression levels (Barzman et al. 2015). A cornerstone tool of IPM is biological control (Eilenberg et al. 2001; Stenberg 2017; Buitenhuis et al. 2023), which focuses on the intrinsic pest control provided by natural enemies and other beneficial organisms (henceforth beneficials). Biological control has a demonstrable trajectory of managing pests, either by conserving or promoting existing populations of beneficials in non-crop habitats (Cullen et al. 2008; Thomson & Hoffmann 2009; Gagic et al. 2018; Alarcón-Segura et al. 2022) or by augmenting beneficial populations through inoculating or inundating the crop with commercially reared individuals (Collier & Van Steenwyk 2004; van Lenteren et al. 2018). A key ecological trait of beneficials that shape their role as biological control agents is their degree of dietary specialisation (Loxdale et al. 2011; Zikic et al. 2017), with some species consistently or opportunistically preying on a broad range of pests (e.g. most ladybirds and spiders), while others show very specific host associations (e.g. some aphid parasitoid wasps). Importantly, resource utilisation and feeding mechanism complementarities and synergies along this specialisation continuum can have positive effects on pest control (Tylianakis & Romo 2010; Gontijo et al. 2015; Jonsson et al. 2017; Alhadidi et al. 2018). Similarly, effective pest control relies on beneficials being active and distributed along seasonal and regional continua, ranging from species that are active during a limited period within a season and endemic to a specific area (Furlong et al. 2004), to those that are globally distributed and active throughout the entire year (Brown et al. 2011).

A recent development in IPM practice has been the emergence of selective pesticides (Umetsu and Shirai 2020; Lykogianni et al. 2021) designed to be effective against target pests while causing minimum negative effects on beneficials. These selective pesticides are increasingly being adopted across agricultural industries and their preference over broad-spectrum chemicals is being encouraged by IPM researchers and practitioners (Gentz et al. 2010; Umina et al. 2015; Torres & Bueno 2018). Some studies have investigated the lethal, sub-lethal and transgenerational effects of selective pesticides on a range of beneficial species (Croft & Whalon 1982; Flexner et al. 1986; Fountain & Medd 2015; Mills et al. 2016; Siviter & Muth 2020; Wu et al. 2022; Knapp et al. 2023a; Overton et al. 2023; McDougall et al. in press). However, evidence on whether these chemicals can control targeted pests without precipitating adverse effects across a range of beneficials including those that are regionally and seasonally restricted remains scarce.

At present, most farmers and advisors considering sustainable chemical choices for resident and augmented beneficials make decisions on toxicity ratings derived from data from a single or a few widespread species rather than locally occurring beneficials. This issue is compounded by decisions that are based on chemicals tested in widely distributed species that are meant, but unlikely, to represent the pool of regionally relevant species potentially acting as biocontrol agents in a given region and season. Importantly, an emerging body of research points to considerable differences in the responses of closely related species to both broad-spectrum and selective chemicals (Bergeron & Schmidt-Jeffris 2020; Duso et al. 2020; Overton et al. 2023). This raises the issue of whether reported chemical toxicities can be generalised across the range of beneficial insects, mites and other invertebrates providing biological control. There are also experimental limitations on past work, including: (1) inconsistencies in testing methods employed, which make it difficult to draw like-for-like comparisons across species; (2) a lack of evaluation of how standardised toxicity ratings may be influenced by experimental decisions around application rates and exposure timepoints; and (3) limited assessments for species or groups of species beyond the range of mass-reared, commercially available species (see Overton et al. 2021).

Here, we report on the acute toxicity response of beneficial insects and arachnids to both broad-spectrum and selective pesticides. We conducted a series of standardised ecotoxicology bioassays to assemble a comprehensive dataset documenting the acute toxicity effects of 20 chemicals (representing 14 mode of action groups, including broad-spectrum and selective pesticides) on 15 beneficial species (representing a broad range of predator and parasitoid groups) at three exposure periods. We then assessed how mortality varied among chemicals, species, and exposure timepoints, aiming to: (1) identify and summarise both common and diverging patterns across the gamut of acute toxicity responses to pesticides displayed by the tested species; (2) determine if the effect of pesticides on beetles, wasps and mites (widely regarded as the most important arthropod biocontrol agents in agriculture) is consistent across the tested species within these groups; and (3) assess how mortality changes across exposure timepoints – therefore influencing standardised toxicity ratings such as those currently employed by the International Organisation for Biological Control (IOBC).

## 2. Material and methods

### 2.1 Experimental approach

#### 2.1.1 Beneficial insects and arachnids

We conducted acute toxicity assessments on 15 beneficial invertebrate species, spanning ten predatory and parasitoid insect and arachnid groups, including predatory mites, spiders, ladybirds, rove beetles, lacewings, hoverflies, predatory bugs, aphid parasitoid wasps, and lepidopteran egg and larval parasitoid wasps (Tables S1 & S2). Although selected because they are important beneficials of the Australian grains industry (see Overton et al. 2021), many of these species are among the most important beneficials globally (Buitenhuis et al. 2023). Local representative study species for each group were informed by a gap analysis conducted by Overton and colleagues (2021) and in consultation with industry experts. These species were predominantly sourced from local commercial suppliers, except hoverflies (*Melangyna spp*.), wolf spiders (*Venatrix spp*.) and snout mites (*Odontoscirus lapidaria*), which were sourced through field collections in Victoria, Australia. Adult hoverflies were collected from flowering plants (*Brassica spp*., *Taraxacum spp*. and *Acacia spp*.) in the spring of 2021. The tested larvae were then obtained by inducing oviposition in the laboratory, which was achieved by providing gravid females with bok choy (*Brassica rapa*) plants infested with green peach aphids (*Myzus persicae*). Snout mites were directly collected during the Austral winter growing seasons of 2022 and 2023 from pastoral fields (Knapp et al. 2023a). Female wolf spiders carrying egg cases or spiderlings were collected between spring and autumn of 2021 and 2022, and spiderlings were tested following dispersal from their mother’s abdomen.

#### 2.1.2 Pesticides

We tested a comprehensive range of pesticide formulations, comprising 14 mode of action groups and 20 active ingredients (Table S3), with each pesticide generally tested at the Maximum Registered Field Rate (MRFR) in Australian grain crops (APVMA 2021). Solutions of each pesticide were made up by diluting the necessary volume of product in deionised water.

#### 2.1.3 Acute mortality bioassays

We examined the acute mortality responses of beneficials to pesticides over multiple laboratory bioassays between 2020-2023. Each bioassay included a negative control (deionised water). The gap analysis by Overton and colleagues (2021) informed the selection of pesticides tested on each beneficial species, with priority given to species x pesticide combinations for which no prior literature was available.

In all bioassays, beneficials were exposed to dried pesticide spray residues in line with IOBC protocols (Sterk et al. 1999), as described in full detail by Knapp and colleagues (2023a) and Overton and colleagues (2023). In brief, a potter spray tower (Burkard Manufacturing, Rickmansworth, United Kingdom) was used to coat 35 mm petri dish bases with even and consistent spray deposits of 1 mg/cm^2^, equivalent to a field application rate of 100 L/ha of diluted pesticide. Residues were allowed to dry in a fume hood for a minimum of 30 min before beneficials were introduced to dishes. We typically tested a minimum of 30 individuals for each species and treatment combination, but this was not always possible, particularly for field-collected species. Parasitoid wasps were housed three per dish, while insect and arachnid predators were housed in individual dishes to prevent cannibalism (Table S3). Dishes were supplied with different food and humidity sources depending on the requirements of each species, along with other minor species-specific methodological differences (Table S3). All dishes were maintained at room temperature (20-25 0C). Individuals were scored as alive (actively moving), dead (no movement) or incapacitated (inhibited movement) and at three timepoints: 24, 48 and, in most cases, 72 h after treatment (HAT).

### 2.2. Dataset management

Our study’s dataset is part of a larger ongoing project as part of the Australian Grains Pest Innovation Program (AGPIP), which produces and curates an ‘Australian grains beneficials chemical toxicity table’ (Knapp et al. 2023b). The project’s empirically derived dataset, currently comprising over 40,500 datapoints, has been previously published and is publicly available (Umina et al. 2024). The final number of datapoints for each chemical and species combination included in our study is provided in Table S4. All incapacitated individuals were pooled with dead individuals for analysis, as they are unlikely to survive or contribute to the next generation (Hoffmann et al. 1997; Roberts et al. 2011).

### 2.3 Statistical modelling

#### 2.3.1 Core mortality model

We used a variation of the binomially distributed generalised linear model described by Kéry (2010) to assess how mortality varied across chemicals and species. The model was specified as:

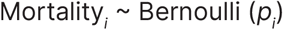

where *p*_*i*_ is the probability that the tested individual in replicate petri dish *i* will be dead. The linear predictor was specified on the logit-probability scale as:

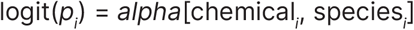

where chemical_*i*_ is an integer indexing the chemical treatments; species_*i*_ an integer indexing the species; and *alpha* the combined chemical/ species treatment-level fix effects for the potential chemical/species combinations, which were specified as two-dimensional matrices and assigned to non-informative Normal priors with mean = 0 and precision = 0.001. In the interest of code simplicity, we modelled data from each of the three HAT timepoints in individual batches. An alternative approach, that would have yielded the same estimates, would be to specify *alpha* as a three-dimensional matrix, with a corresponding hat_*i*_ integer indexing the three HAT timepoints.

We ran data from each bioassay through this core model, using the following variation of the method proposed by Abbott (1925) to control for baseline mortality (i.e. the mortality observed in the water negative control):

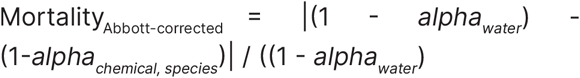

As these calculations were conducted within a Bayesian inference framework, the Abbott-corrected mortality estimates were derived with their full associated uncertainties.

We used the posterior estimates to map the probability or probabilities that the mortality response of a species will be classified into one or more of the IOBC toxicity classifications, where 1 = harmless (<30% mortality), 2 = slightly harmful (30-79% mortality), 3 = moderately harmful (80-99% mortality), and 4 = harmful (>99% mortality).

#### 2.3.2 Mortality by chemical and mortality by species meta-analyses

We used a variation of the meta-analytical approach described by Palma and colleagues (2016) to assess how mortality varied across chemicals and how mortality varied across species. Specifically, we used the mean and standard deviation mortality estimates derived from the 48-HAT models to parameterise 35 meta-analyses (one for each chemical and species). Each meta-analysis was specified as:

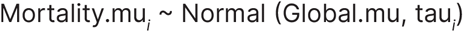

where the global mean Global.mu was given a Normal (0, 0.0001)T(0,1) prior and the precision was specified as:

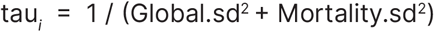

where the global standard deviation Global.sd was given a Uniform (0, 100) prior.

For the mortality by species meta-analyses, we used the approach described in 2.3.1 to map the probability or probabilities that the mortality response of species will be grouped into one or more of the IOBC toxicity classifications. We selected 48-HAT data to ensure standardisation across all species in our study and because it is the most common exposure period used to test acute toxicity effects across a broad range of beneficials globally.

#### 2.3.3 Differences in IOBC classification by beneficial group

We used a Bernoulli distributed generalised linear model to determine if the effect of pesticides on beetles (*Hippodamia variegata, Harmonia conformis, Dalotia coriaria*), wasps (*Aphelinus abdominalis, Aphidius colemani, Diaeretiella rapae, Diadegma semiclausum, Trichogramma pretiosum*) and mites (*Hypoaspis aculeifer, Typhlodromus montdorensis, Odontoscirus lapidaria*) was consistent across the tested species within these groups. Specifically, we modelled the probability that the mortality of species representing a given beneficial group fall into more than one IOBC classification. This assessment was limited to the 48-HAT timepoint and to those beneficial groups in our dataset with three or more representative species (i.e. beetles, wasps and mites). We developed an algorithm that inspected the mean mortality estimates from the chemical/species models, assigned each species to its corresponding IOBC classification, and, for chemicals represented by two or more species, evaluated the resulting IOBC assignations, coding a 0 when the species were consistently mapped onto a single IOBC classification and 1 when they were mapped onto more than one. This vector was then modelled as:

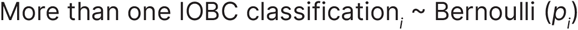

where *p*_*i*_ is the probability that the species mortalities within the beneficial group in replicate chemical *i* map onto more than one IOBC classification. The linear predictor was specified on the logit-probability scale as:

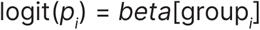

where group_*i*_ is an integer indexing the beneficial group treatments (1 = beetles, 2 = wasps, and 3 = mites); and *beta* the beneficial group treatment-level fix effects, which were assigned to non-informative Normal priors with mean = 0 and precision = 0.001.

#### 2.3.4 Change in IOBC classification by exposure period

We used a Bernoulli distributed generalised linear model to determine if the effect of pesticides on species is consistent across the tested HAT exposure timepoints. Specifically, we modelled the probability that the IOBC classification will transition from a lower to a higher level as mortality is evaluated at different HAT timepoints. We developed an algorithm that inspected the mean mortality estimates from the chemical/species/ HAT models (Tables S5, S6 & S7), assigned each species to its corresponding IOBC classification, and evaluated the resulting IOBC assignations, coding a 0 when the IOBC classification did not experience a level transition and 1 when it did. This vector was then modelled as:

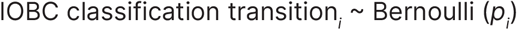

where *p*_*i*_ is the probability that the IOBC classification in replicate chemical/species i will transition from a lower to a higher classification. The linear predictor was specified on the logit-probability scale as:

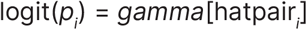

where hatpair_*i*_ is an integer indexing the HAT transition pair treatments (1 = 24-HAT to 48-HAT, 2 = 48-HAT to 72-HAT, and 3 = 24-HAT to 72-HAT); and *gamma* the HAT transition pair treatment-level fix effects, which were assigned to non-informative Normal priors with mean = 0 and precision = 0.001.

#### 2.3.5 Bayesian inference implementation

We estimated model parameters under Bayesian inference, using Markov Chain Monte Carlo simulations to draw samples from the parameters’ posterior distributions. Models were coded and run in JAGS (Plummer 2003), which was accessed through the R package jagsUI (Kellner 2016). We used three chains of 5,000 iterations, discarding the first 500 in each chain as burn-in. To confirm acceptable convergence levels, we visually inspected the chains and verified that the values of the Gelman-Rubin statistic were lower than 1.1 (Gelman and Hill 2007).

#### 2.3.6 Pairwise mean group effect comparisons

To assist with interpretation of modelling outputs, we ran the posterior estimates from the core mortality models and the mortality by chemical and mortality by species meta-analyses through the pairwise mean group effect comparison algorithm described by Knapp and colleagues (2023a). The algorithm generates pairwise mean group effect triangular tables by first comparing the estimated 95% credible intervals (CI) of each group pair and then: (1) estimating the multiplicative group effect separating the smaller effect from the larger, when the group pair’s CIs do not overlap; or (2) labelling the pairwise comparison as ‘not statistically different’, when the CIs overlap.

## 3. Results

### 3.1 Mortality by chemicals meta-analyses

Mortality varied extensively across chemicals (Figure 1; Table S5). Responses ranged from chemicals that showed very high acute toxicity effects (mean mortality > 99.5%) such as the carbamate methomyl and the organophosphates chlorpyrifos and dimethoate – which fall into the IOBC classification 4 (harmful) with over 98.5% certainty – to chemicals such as chlorantraniliprole, *Bacillus thuringiensis* (*Bt*), and Nuclearpolyhendrosis virus (NPV) that showed very low toxic effects (mean mortality < 1.5%) – which fall into the IOBC classification 1 (harmless) with 100% certainty (Figure 1; Table S5). The overall effects of the most toxic two chemicals (chlorpyrifos and dimethoate) were statistically different, and on average, approximately 50 times greater than that showed by the least toxic chemical (chlorantraniliprole) (Table S6).

**Figure 1.**
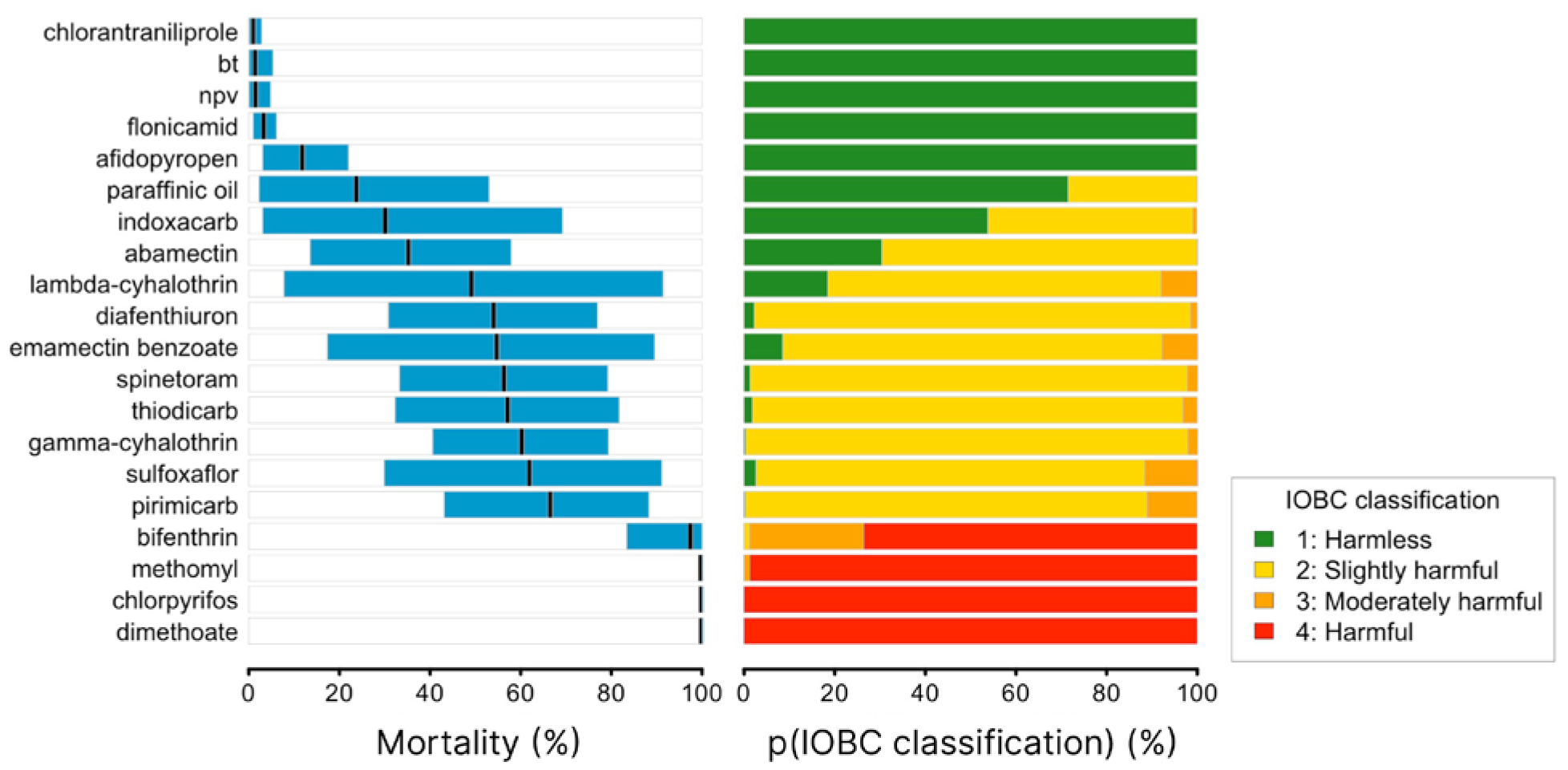
Mortality responses of beneficial insect and arachnid species to each of the tested chemicals at 48-HAT. Black lines indicate the mean response and the blue rectangles the associated statistical uncertainty (95% CIs). The right-hand panel shows the probability, or probabilities, that the mortality response to a given chemical treatment falls into one or more IOBC toxicity classifications, where 1 = ‘harmless’ (< 30% mortality; green), 2 = ‘slightly harmful’ (30-79% mortality; yellow), 3 = ‘moderately harmful’ (80-99% mortality; orange), and 4 = ‘harmful’ (> 99% mortality, red). CIs: credible intervals; IOBC: International Organisation for Biological Control; HAT: hours after treatment; p: probability.

### 3.2 Mortality by species meta-analyses

Mortality also varied considerably across species; however, no species showed the very high or very low acute toxicity effects observed in the mortality by chemical responses (Figure 2; Tables S7 & S8). Those species experiencing the greatest acute toxicity effects, across all chemicals tested, were the large spotted ladybird *Harmonia conformis* and the lepidopteran egg parasitoid wasp *Trichogramma pretiosum* (mean mortality > 74.5%; Figure 2; Table S7). The effects on these species were statistically different, and on average, approximately three times greater than the toxicity effects experienced by hoverflies (*Melangyna spp*.) and the rove beetle *Dalotia coriaria* (Table S8), which experienced the lowest toxicity effects (mean mortality < 25.5%; Figure 2; Table S7).

**Figure 2.**
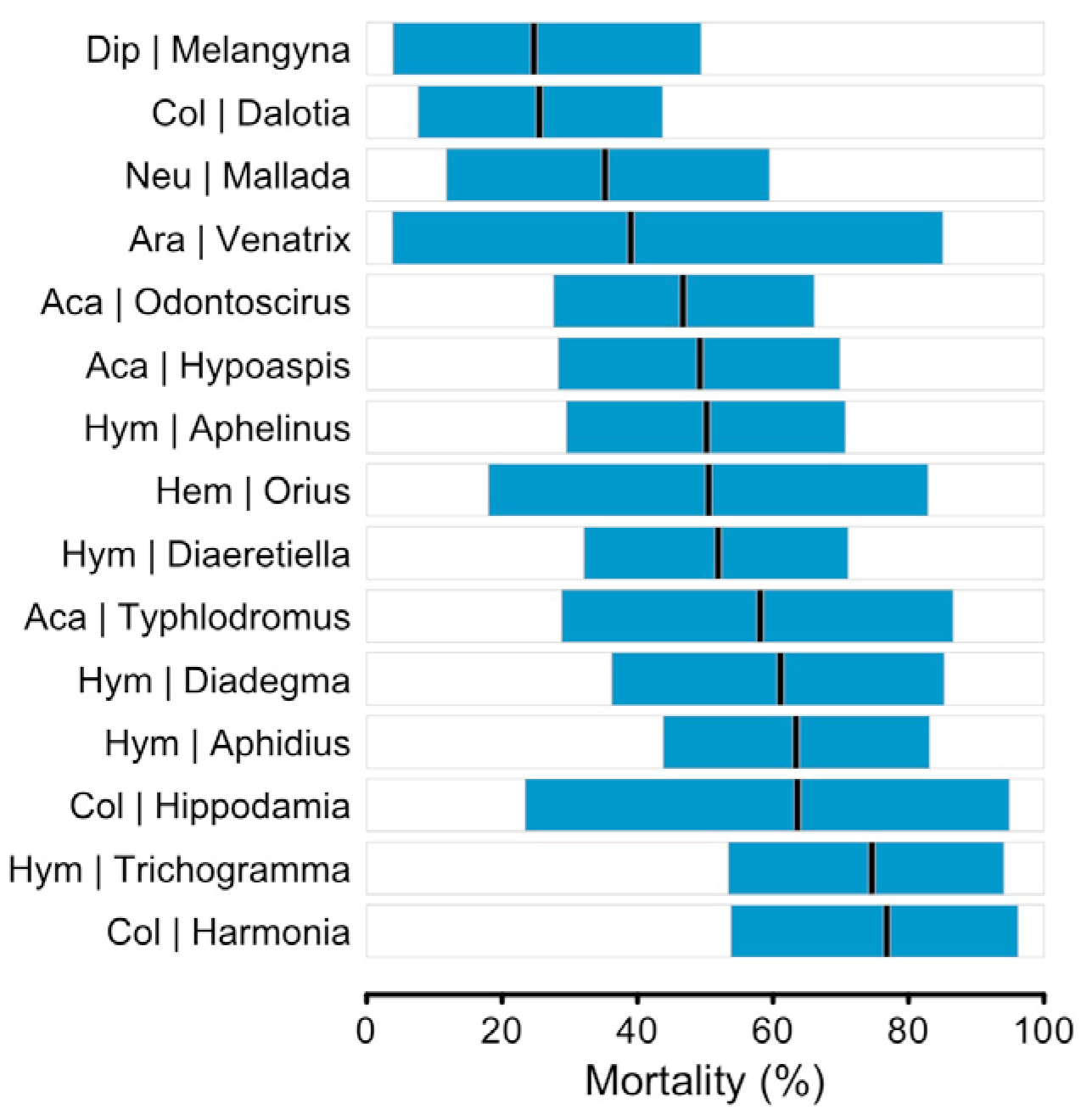
Mortality responses of beneficial insect and arachnid species across all chemicals at 48-HAT. Black lines indicate the mean response and the blue rectangles the associated statistical uncertainty (95% CIs). CIs: credible intervals.

#### 3.3 Mortality by chemicals and species analyses

The effect of chemicals on mortality across species showed two distinct patterns. On the one hand, some chemicals showed either very high or very low responses that were highly or fully consistent across species, meaning the chemical fall into a single IOBC classification (e.g. classification 1: Bt, chlorantraniliprole and NPV; classification 4: methomyl and chlorpyrifos; Figure 3; Table S9). On the other hand, a range of chemicals showed responses that varied across species – sometimes moderately (e.g. flonicamid) but often substantially (e.g. spinetoram) – which prevented the chemical from being confidently mapped into a single IOBC classification (Figure 3; Table S9). Among this later group were both broad-spectrum (e.g. the synthetic pyrethroid gamma-cyhalothrin) and selective (e.g. diafenthiuron) chemicals that showed species-specific responses that fell within all four IOBC classifications (Figure 3; Table S9). In both cases, these chemicals showed either a bimodal response – where species aggregated into a largely IOBC classification 1 group and a largely IOBC classification 4 group – or a continuous response – where the toxicity responses of species span the full mortality gradient (Figure 3). An example of the bimodal response was sulfoxaflor, in which the effects on the predominantly group 4 classification were statistically different, and on average, between approximately 4 to 23 times greater than the effects on the largely group 1 classification (Table S10). The continuous response is exemplified by spinetoram, which included species with effects that were statistically different to each other across all four IOBC classifications; for instance, the effect on the aphid parasitoid wasp *Diaeretiella rapae* (IOBC classification 4) was 1.1 times greater than the effect on the lepidopteran egg parasitoid wasp *Trichogramma pretiosum* (IOBC classification 3 - ‘moderately harmful’), 1.7 times greater than the effect on the large spotted ladybird *Harmonia conformis* (IOBC classification 2 - ‘slightly harmful’), and 23.2 times greater than the effect on the snout mite *Odontoscirus lapidaria* (IOBC classification 1) (Figure 3; Table S10).

**Figure 3.**
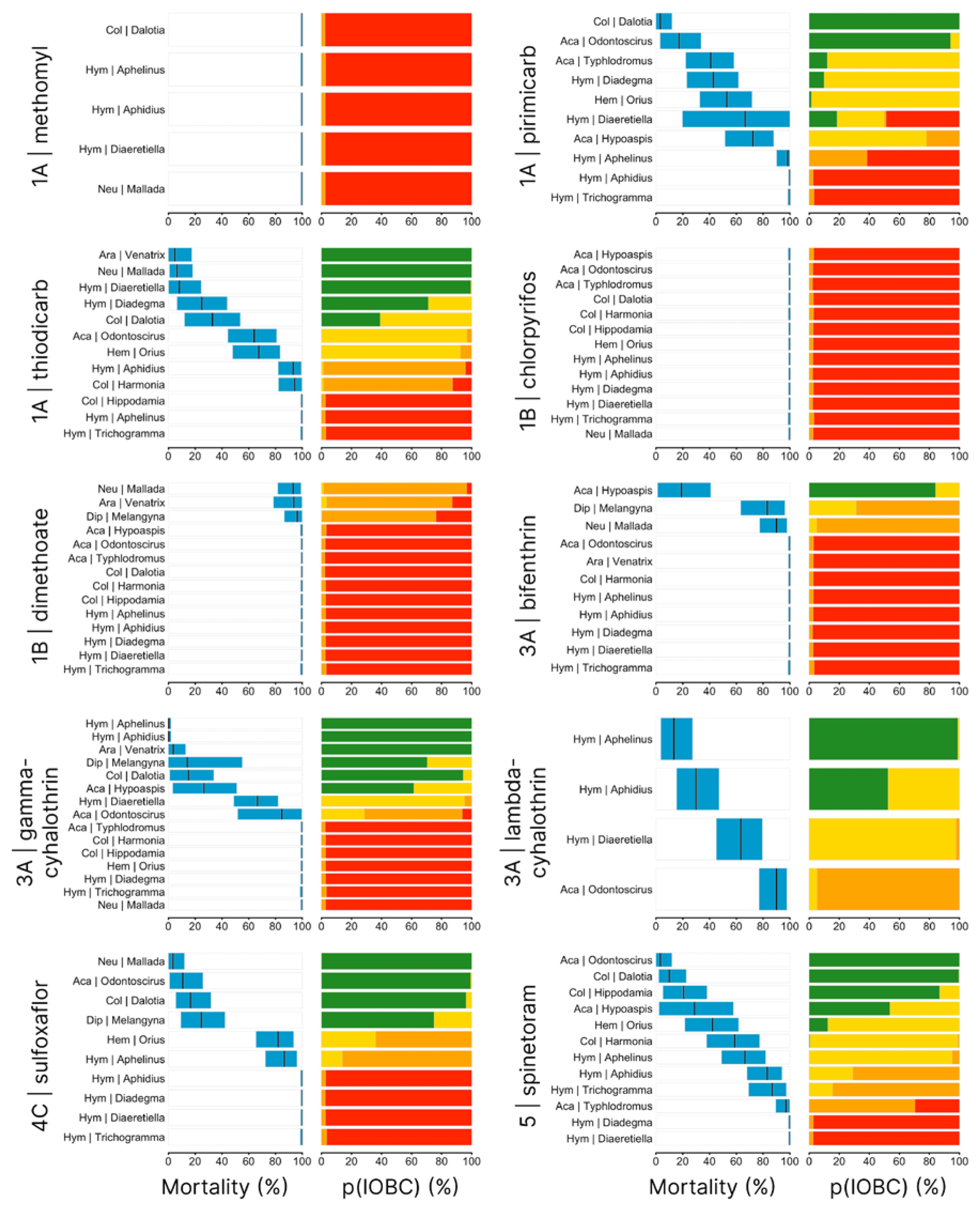
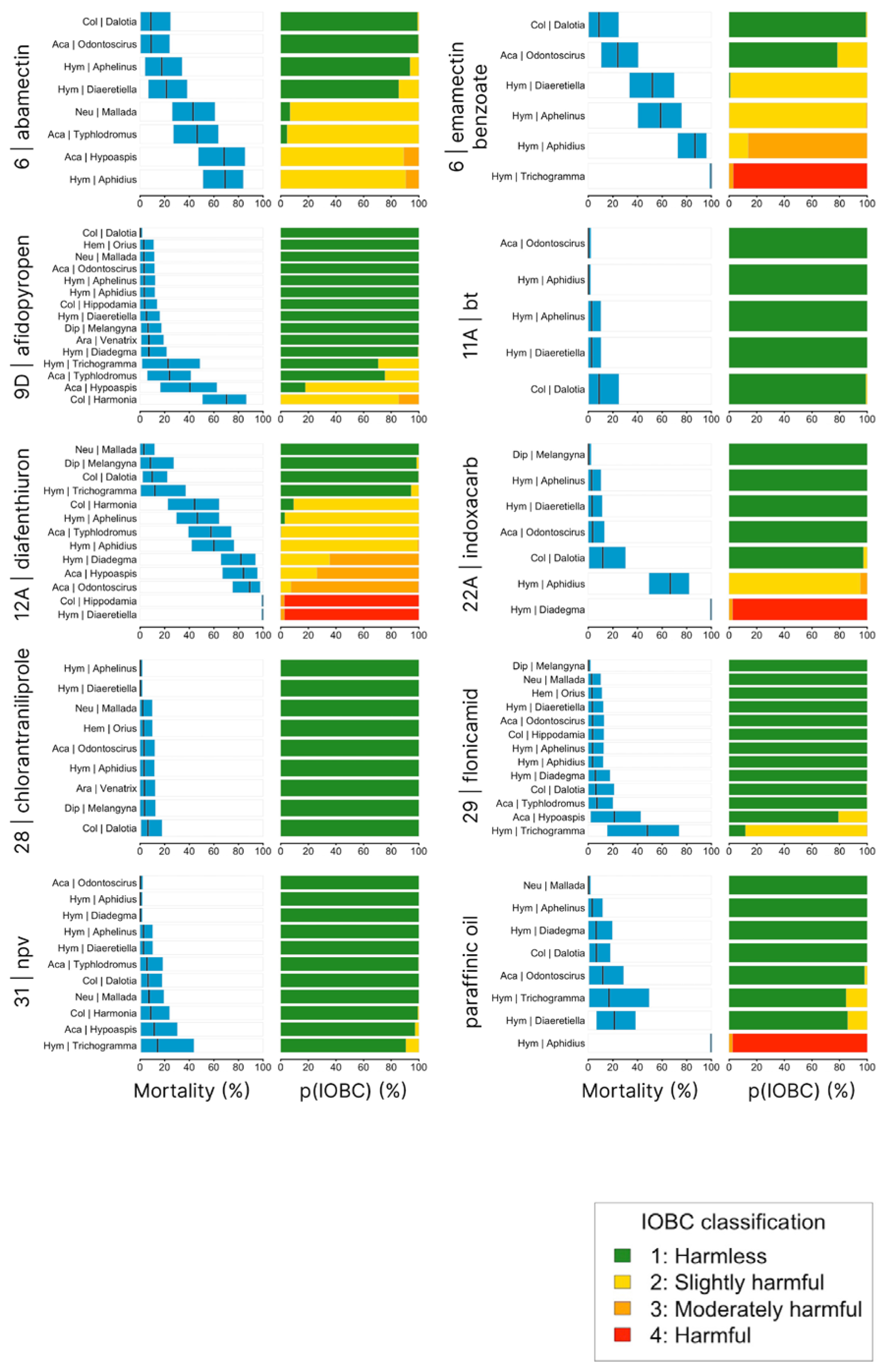
Species-specific mortality responses of beneficial insect and arachnid species to each of the tested chemicals at 48-HAT. Black lines indicate the mean response and the blue rectangles the associated statistical uncertainty (95% CIs). The right-hand panel shows the probability, or probabilities, that the mortality response to a given chemical treatment falls into one or more IOBC toxicity classifications, where 1 = ‘harmless’ (< 30% mortality; green), 2 = ‘slightly harmful’ (30-79% mortality; yellow), 3 = ‘moderately harmful’ (80-99% mortality; orange), and 4 = ‘harmful’ (> 99% mortality, red). CIs: credible intervals; IOBC: International Organisation for Biological Control; HAT: hours after treatment; p: probability.

### 3.4 Differences in IOBC classification by beneficial group

The probability that species within a beneficial group across all chemicals fall into more than one IOBC classification was, on average, higher for wasps (60%) and mites (58%) than for beetles (50%), however these were not statistically different to one another (Table S11). For beetles and mites, most chemicals that showed different responses involved two adjacent IOBC classifications (Figure 4; Table S12). The most frequent difference in these groups was from classification 1 to classification 2, while some broad-spectrum chemicals showed more pronounced differences, varying from classification 1 to classification 4 (Figure 4; Table S12). For wasps, most chemicals that showed different responses involved three IOBC classifications, mostly from classification 2 to classification 4 (Figure 4; Table S12). In the case of diafenthiuron, differences in toxicity effects between wasp species spanned all four IOBC classifications (Figure 4; Table S12).

**Figure 4.**
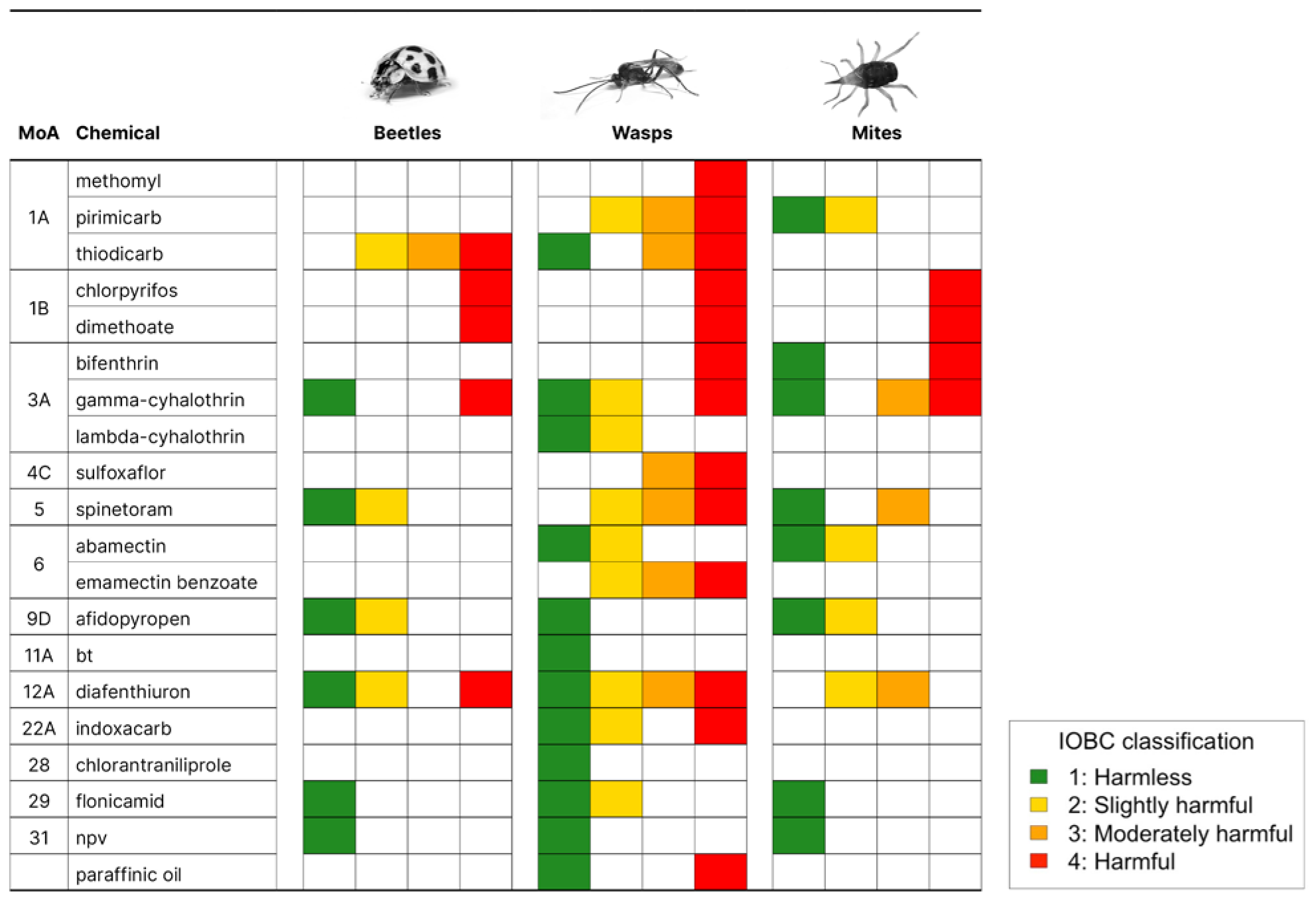
IOBC toxicity classifications by chemical and beneficial group. IOBC classifications are 1 = ‘harmless’ (< 30% mortality; green), 2 = ‘slightly harmful’ (30-79% mortality; yellow), 3 = ‘moderately harmful’ (80-99% mortality; orange), and 4 = ‘harmful’ (> 99% mortality, red). IOBC: International Organisation for Biological Control; MoA: mode of action.

### 3.5 Change in IOBC classification by exposure period

The probability of a change in IOBC classification was on average relatively low when mortality was assessed at 48-HAT compared with 24-HAT (24%) and at 72-HAT compared with 48-HAT (18%) but increased to 34% when assessed at 72-HAT compared with 24-HAT (Table S13). When transitions between IOBC classifications were documented, most 24-to 48-HAT and 48- to 72-HAT shifts were between two adjacent IOBC classifications, while 24-to 72-HAT shifts included several transitions across three or even four IOBC classifications (Table S14). Examples of the latter shifts included the effect of afidopyropen on the large spotted ladybird *Harmonia conformis* – which was on average 3.3 times greater at 72-HAT compared with 24-HAT, shifting its IOBC classification from 1 to 3 – and the effect of diafenthiuron on the lepidopteran parasitoid *Diadegma semiclausum* – which was on average 18.9 times greater at 72-HAT compared with 24-HAT, shifting its IOBC classification from 1 to 4 (Table S14).

## 4. Discussion

In this study, we provide multiple threads of evidence for both strong consistencies and large variations in the acute toxicity response of beneficial insects and arachnids to broad-spectrum and selective pesticides. Our results also indicate that the effect of pesticides on beetles, wasps and mites is not consistent across the tested species within these groups, with the toxicity of most chemicals falling into two or more IOBC classifications depending on the beneficial species tested. Finally, we show that the acute toxicity effects of chemicals on beneficials increase across experimental exposure timepoints, influencing the standardised toxicity ratings currently employed by the IOBC. Taken together, our findings demonstrate that in most cases, chemical toxicities cannot be generalised across a range of beneficial insects and mites providing biological control. This empirical evidence, particularly if supplemented with future field-based studies, will support farmers and their advisors endeavouring to manage crop pests in a way that minimises the undesirable impacts on beneficial species.

### 4.1 Species-specific effects of broad-spectrum and selective chemicals

Our results indicate that the acute toxicity effects of broad-spectrum and selective chemicals vary extensively across beneficial species. Not unexpectedly, some broad-spectrum (e.g. the carbamate methomyl and the organophosphates chlorpyrifos and dimethoate) and selective (e.g. the diamide chlorantraniliprole, the baculoviruses NPV and the bacterium *Bt*) chemicals showed high and low acute toxicity effects, respectively, that were fully consistent across all beneficials that we tested. This consistency of responses across multiple beneficial species is congruent with numerous other laboratory and field studies with these chemicals (Jepson et al. 1995; Armenta et al. 2003; Federici 2003; Suma et al. 2009; Brugger et al. 2010; Liu et al. 2016; Hill et al. 2017).

A striking result to emerge, however, is that some of the other broad-spectrum and selective chemicals we tested showed responses that varied depending on the beneficial species on which they were tested. Notable examples among the broad-spectrums included the carbamates pirimicarb and thiodicarb, and the synthetic pyrethroids bifenthrin and gamma-cyhalothrin, while the species-dependent pattern among the selectives was exemplified by the sulfoximine sulfoxaflor, the spinosyn spinetoram, the avermectin emamectin benzoate, the pyropene afidopyropen, and diafenthiuron. Large differences in responses between species (including closely related species) has been observed previously (e.g. Bergeron & Schmidt-Jeffris 2020; Duso et al. 2020), although our results suggest that this phenomenon may be more widespread than previously appreciated. This could easily lead to situations in the field where so-called selective chemicals are just as harmful to locally relevant beneficials as the ‘broad-spectrums’.

### 4.2 Effects of chemicals on beetles, wasps and mites

We found strong evidence that the effect of both broad-spectrum and selective pesticides on beetles, wasps and mites is not consistent across the tested species within these groups. For all three groups, the toxicity of most chemicals fall onto two or more IOBC classifications depending on the beneficial species tested, and in one case (diafenthiuron), the differences involved all four IOBC classifications. This shows the large variability in chemical sensitivity among arthropod species, as previously demonstrated by Bergeron & Schmidt-Jeffris (2020) and discussed by Duso and colleagues (2020), and highlights the risk of making generalised toxicity inferences based on data from a single (or small number of) species. That being said, some chemicals showed consistent responses within the three beneficial groups examined in this study (e.g. chlorpyrifos, dimethoate, chlorantraniliprole, *Bt*, NPV), something other studies have also shown. For example, Jepson and colleagues (1995) reported consistent harmful effects of the organophosphate dimethoate on six predatory beetles, while Brugger and colleagues (2010) reported consistent harmless effects of the diamide chlorantraniliprole on seven species of parasitoid wasps, including many of the species and genera included in our study.

### 4.3 Scoring timepoint decisions influence IOBC toxicity classifications

This work shows that key experimental decisions, such as whether to assess mortality at 24-, 48- or 72-HAT, can influence the outcomes of standardised toxicity ratings (such as the IOBC classifications). In one third of cases, chemicals transitioned from lower to higher IOBC classifications when assessed at 72-HAT compared with 24-HAT, including cases where a chemical fully transitioned from harmless to harmful (e.g. diafenthiuron on the lepidopteran parasitoid *Diadegma semiclausum*). Such findings are not surprising and have been shown previously (e.g. Lima et al. 2016) yet remain important to highlight. They point to limitations of IOBC protocols and suggest that for some species (e.g. parasitoid wasps - which are ordinarily assessed after 48-HAT) mortality assessments at later exposure time points would be useful.

### 4.4. Limitations and future research

Our standardised approach to quantify the acute toxicity effects of chemicals on beneficials helps overcome previous experimental limitations, allowing for robust like-for-like comparisons across species, including field-collected species not available through mass-rearing commercial insectaries. However, there are numerous limitations that could be improved in future studies. Time and resource restrictions did not allow us to (1) test all combinations across the chemical/species we worked with, (2) test multiple populations of the same species, or (3) incorporate other chemicals – across a wider range of mode of actions – and other species – across a wider range of beneficial groups (e.g. pollinators, detritivores). Research by our group is already underway to address some of these issues, with an emphasis on further understanding the effects on non-commercially available species such as spiders and effects caused by other increasingly used selective pesticides. On a wider level, our work focused on pesticides, and hence did not incorporate herbicides or fungicides. Although arguably less important than pesticides in this field of research, herbicides and fungicides can also lead to non-target effects on beneficials (Jansen 1999).

Our study only investigated acute toxicity effects, and while valuable, there is a growing body of empirical work that shows both broad-spectrum and selective pesticides can have sub-lethal and transgenerational effects on beneficials at field-realistic levels of exposure (Desneux et al. 2007; Mills et al. 2016; Siviter & Muth 2020; Wu et al. 2022; McDougall et al. in press). These investigations – spanning work on development, longevity, fecundity, mobility, orientation, feeding, oviposition, learning, metabolism and detoxification, among many other key physiological, behavioural and genetic effects – highlight how acute toxicity effects estimated from studies such as ours only partially reveal the gamut of potentially deleterious effects of pesticides on beneficials. As others before us (e.g. Desneux et al. 2007), we recommend that efforts to robustly understand, and in time successfully address, the true range of detrimental effects of pesticides on beneficials should complement acute toxicity experimentation, with the more nuanced perspective provided by sub-lethal, transgenerational and gene expression investigations; an approach that is well-aligned with current IOBC practice guidelines (Hassan 1989).

Another important focus of future studies should be validation of laboratory-based findings under greenhouse, semi-field and field conditions. The consistency of chemical toxicity effects across experimental scales is a nuanced and complex process (Macfadyen et al. 2014; Overton et al. 2021), particularly given the diverse array of different crop types and environmental and crop growing conditions. If our study was conducted under more representative conditions found in the field, toxicities may vary from those observed here. Nonetheless, we believe multi-chemical, multi-species synthesis laboratory studies, such as the one we report here, are valuable for benchmarking baseline acute toxicity effects.

Finally, there are several questions that warrant further attention. For example, what makes certain pesticide groups consistently toxic or non-toxic against a diverse range of beneficials? Does this align with expectations based on their insecticidal/arachnicide mechanisms? While much is already known about the selectivity of biological pesticides, such as *Bt* and NPV (Armenta et al. 2003; Federici 2003), this is less clear for many synthetic compounds. Whether this line of inquiry could help inform the development of new chemistries that are truly harmless to non-target beneficials remains to be seen.

## Acknowledgements

This research was funded by the Grains Research and Development Corporation, grant number UOM1906–002RTX. We thank Bugs for Bugs and Biological Services for providing insect and mite samples and those agrichemical companies that provided pesticide samples. Special thanks to Aston Arthur, Xuan Cheng, Evatt Chirgwin, Lisa Kirkland, Karyn Moore, Esti Palma, Olivia Reynolds, Julia Severi and Samantha Ward for their contributions to this research. The authors would like to acknowledge the Traditional Custodians of the land and waterways on which this research took place, the Woi wurrung and Boon wurrung peoples of the eastern Kulin Nations. We pay our respects to their Elders, past, present, and emerging.

## CRediT author statement

**Luis Mata**: Formal analysis, Data curation, Writing - Original Draft, Visualisation, Supervision. **Rosemary Knapp**: Investigation, Data curation, Writing - Review & Editing, Visualisation. **Robert McDougall**: Investigation, Writing - Review & Editing. **Kathy Overton**: Methodology, Investigation, Writing - Review & Editing. **Ary Hoffmann**: Writing - Review & Editing, Funding acquisition. **Paul Umina**: Conceptualisation, Methodology, Investigation, Writing - Review & Editing, Supervision, Project administration, Funding acquisition.

## Data availability statement

Data and codes to reproduce models, plots and tables are already published and publicly available in Zenodo. Data: https://doi.org/10.5281/zenodo.8368898 (Umina et al. 2024). Codes: https://doi.org/10.5281/zenodo.10570361 (Mata 2024).

